# HSPA8 chaperone complex drives chaperone-mediated autophagy regulation in acute promyelocytic leukemia differentiation

**DOI:** 10.1101/2022.08.07.502745

**Authors:** Sreoshee Rafiq, Irene Mungure, Nicolas J. Niklaus, Stefan Müller, Arnaud Jacquel, Guillaume Robert, Patrick Auberger, Bruce E. Torbett, Sylviane Muller, Mario P. Tschan, Magali Humbert

## Abstract

Acute myeloid leukemia (AML) is a cancer of the hematopoietic system characterized by the hyperproliferation of undifferentiated cells of the myeloid lineage. While most of AML therapy are focusing on tumor debulking, all-*trans* retinoic acid (ATRA) induces differentiation in acute promeylocytic leukemia (APL) a particular subtype. Macroautophagy has been extensively investigated in the context of various cancers and is often dysregulated in AML where it can have context-dependent pro- or anti-leukemogenic effects. On the contrary, the implications of chaperone-mediated autophagy (CMA) on the pathophysiology of diseases are still being explored and its role in AML has remained elusive. To answer our questions we took advantages of human AML primary samples and databases. Furthermore, we used ATRA-sensitive (NB4) and –resistant (NB4-R1) cells to further dissect a potential function for CMA in ATRA-mediated neutrophil differentiation. NB4-R1 cells are unique in that they do respond to retinoic acid transcriptionally, but do not mature in response to retinoid signaling alone unless maturation is triggered by adding cAMP. Here, we report that CMA related mRNA transcripts are higher expressed in immature hematopoietic cells as compared to neutrophils. In line, lysosomal degradation of mCherry-KFERQ CMA reporter decreases during ATRA-induced differentiation of APL cells. On the other hand, using NB4-R1 cells we found that macroautophagy flux primed the ATRA resistant NB4-R1 cells to differentiate upon ATRA treatment, but reduced association of LAMP-2A and HSPA8 is necessary for complete neutrophil maturation. Accordingly, depletion of HSPA8 attenuated CMA activity and facilitated APL differentiation. In contrast, maintaining high CMA activity by ectopic expression of LAMP-2A impeded APL differentiation. Overall, our findings demonstrate that both normal and APL neutrophil differentiation require CMA downregulation and this pathway is predominantly dependent on HSPA8 assisted by other co-chaperones.

## 1 Introduction

Hematopoiesis is a tightly regulated process that enables hematopoietic stem cells (HSC) to mature into functional hematopoietic cells. Regulated formation of myeloid leukocytes, known as myelopoiesis, is a process paralleled by a reduction in cell growth compared to common myeloid progenitor cells and production of mature myeloid cells with entirely new morphologies and functions (Berliner 2008). Such striking changes in cell architecture require massive remodeling and necessitate maintenance of a delicate balance between macromolecule synthesis and degradation. Therefore, autophagy is highly involved in myeloid cell specific functions, such as monocyte-macrophage phagocytosis and differentiation (Gutierrez et al. 2004; Sanjuan et al. 2007; Jacquel et al. 2012; Zhang et al. 2009), antigen presentation by dendritic cells (Levine and Deretic 2007), and HSC maintenance (Warr et al. 2013; Liu et al. 2007; Mortensen et al. 2011). Due to its essentiality in hematopoiesis, dysregulated autophagy can lead to myeloid malignancies ^10^.

Autophagy is a lysosomal degradation and recycling mechanism of cytosolic components, such as protein aggregates and defective organelles. Three main autophagy pathways have been described: Macroautophagy, chaperone-mediated autophagy (CMA) and microautophagy, the first two being the best studied. Macroautophagy is characterized by the formation of double-membraned vesicles named autophagosomes, which engulf cytoplasmic contents and deliver them to lysosomes (Tanida 2011). In contrast to macroautophagy, CMA cargo delivery does not require the formation of vesicles (Dice 2007). It is a selective autophagy pathway in which the heat shock 70 kDa protein 8 (HSPA8), also known as <u>h</u>eat <u>s</u>hock <u>c</u>ognate protein 70 (HSC70) targets potential client proteins harboring the pentapeptide motif KFERQ (Lys-Phe-Glu-Arg-Gln) to the lysosomal surface where they bind to the lysosome-associated membrane protein (LAMP) type 2A receptor on the lysosomal membrane. LAMP-2A multimers bind to the client-chaperone complex delivering them into lysosomes for subsequent degradation by lysosomal proteases ^13,14^.

Basal macroautophagy is crucial for maintenance of cellular homeostasis in quiescent and proliferating cells ^15^. Several studies have indicated a role of macroautophagy in mammalian development ^16^ and cellular differentiation particularly in adult HSCs and myeloid progenitor cells ^17,18^. Attenuation of macroautophagy results in genomic instability due to events such as, aberrant reactive oxygen species (ROS) production. The tumor-suppressive function of macroautophagy is further supported by animal models, for example heterozygous deletion of *Becn1* in mice displays increased tumor incidence (Qu et al. 2003). Conversely, cancer therapy induced macroautophagy frequently supports tumor cell survival, suggesting an oncogenic function (Hoare, Young, and Narita 2011). Altogether, the role of macroautophagy in tumorigenesis is contentious and depends on the type of tumor and the stage of disease progression.

The AML subtype, acute promyelocytic leukemia (APL), is characterized by the fusion of a key regulator of myeloid differentiation, retinoic acid receptor alpha (RARA) with promyelocytic leukemia (PML) protein, forming the fusion protein PML-RARA which deregulates retinoid signaling ^21^. APL is treated using all-*trans* retinoic acid (ATRA) in combination with arsenic trioxide (ATO), which restore differentiation by activating proteasomal and macroautophagic degradation of PML-RARA aggregates. This further demonstrates the involvement of macroautophagy in myeloid leukemia differentiation ^22,23^. Interestingly, CMA is attenuated by RARA signaling in mouse fibroblasts ^24^ and a frequent crosstalk is observed between CMA and macroautophagy (Massey, Kaushik, and Cuervo 2006; Kaushik et al. 2008). In many cancer cells CMA is upregulated ^27,28^ although similar to macroautophagy, its exact role can be debated. On one hand, CMA degrades the late stage glycolytic enzyme pyruvate kinase M2 (PKM2) which results in accumulation of glycolytic intermediates providing biosynthetic materials for cancer cell growth (Lv et al. 2011). On the other hand, the early stage glycolytic enzyme, hexokinase-2 (HK2) is also a CMA client whose degradation results in a metabolic catastrophe in cancer cells leading to cell death, thereby supporting CMA upregulation as a possible cancer therapy strategy (Xia et al. 2015). CMA also maintains functional HSCs by active participation in lipid metabolism during cellular reprogramming ^31^. Unfortunately, studies on the role of CMA in hematological malignancies such as AML are rather limited and focus mostly on steady state CMA level as opposed to dynamic CMA activity ^32^.

In the present study, we established that reduced expression of CMA related genes allows normal and leukemic granulocytic differentiation. In addition, we measured CMA flux in HSPA8 depleted model and showed that HSPA8 chaperone complex is the limiting factor in CMA activity during APL granulocytic differentiation. Furthermore, we confirmed that reduced HSPA8 facilitates ATRA-mediated APL granulocytic differentiation.

## 2 Materials and methods

### 2.1 Cell lines and primary cell cultures

The human APL cell lines HT93 and NB4 were obtained from American Type Culture Collection and Cell Lines Service (CLS GmBH, Eppelheim, Germany), respectively. Both cell lines as well as ATRA-resistant NB4-R1 cells were maintained in RPMI-1640 (Sigma-Aldrich, Germany) supplemented with 10% (v/v) fetal calf serum (FCS) in a 5% CO_2_-95% air humidified atmosphere at 37°C. 293T cells were cultured in DMEM (Sigma-Aldrich) supplemented with 5% FCS in 7.5% CO_2_ at 37°C. CD34^+^ cells were isolated from cord blood and cultured as previously described ^33^.

### 2.2 Immunofluorescence microscopy

NB4 and HT93 cells were prepared as previously described (Brigger, Proikas-Cezanne, and Tschan 2014). Briefly, cells were fixed with ice-cold 100% (v/v) methanol for 4□min and then incubated with anti-LC3B (3868; Cell Signaling Technology, MA), anti-LAMP-2A (ab18528; Abcam, UK), or anti-HSC70 (MA3-014; Thermo Fisher Scientific, MA) antibodies diluted (1:100) in phosphate-buffered saline (PBS) pH 7.4 containing 1% (w/v) bovine serum albumin (BSA) for 1 h in a humid chamber at room temperature (RT) followed by washing steps with PBS containing 0.1% (v/v) Tween (PBS-T). Then, cells were incubated with the secondary antibodies (cyanine-3 (Cy-3)-labeled anti-rabbit, 111-166-045; fluorescein isothiocyanate (FITC)-anti-rabbit, 111-096-045; Cy3-anti-mouse, 115-166-062; Jackson ImmunoResearch, PA) for 1h at RT. Following washing steps, SlowFade™ gold antifade mountant containing 4′,6-diamidino-2-phenylindole (DAPI) (Thermo Fisher Scientific) was applied to stain nuclei and reduce photobleaching. Images were taken using an Olympus FluoView-1000 (Olympus, Volketswil, Switzerland) confocal microscope at 63x magnification and analyzed using ImageJ software. The plugins RUBEN and JACoP were used for analyzing dot formation and co-localization, respectively.

### 2.3 Cell lysate preparation and western blotting

Whole cell extracts of NB4 and HT93 were prepared using urea lysis buffer and 30-60μg total protein was loaded on a denaturing 12%-polyacrylamide gel self-cast gel (Bio-Rad Laboratories, CA). Proteins were transferred to polyvinylidene fluoride (PVDF) membranes and blocked with 5% (w/v) milk in Tris-buffered saline (TBS) with 0.1% Tween-20 (TBS-T). Blots were incubated with the primary antibodies diluted in blocking solution overnight at 4°C. Following washes with TBS-T, they were incubated with horseradish peroxidase (HRP)-coupled secondary goat anti-rabbit and goat anti-mouse (Cell signaling) at 1:5,000–10,000 for 1h at RT. Primary antibodies used were anti-DAPK2 (2323, ProSci, Poway, CA), anti-ATG5 (#2630, Cell Signaling Technology), anti-LC3B (NB600-1384, Novus Biologicals, Zug, Switzerland), anti-LAMP2A (ab125068, Abcam), anti-HSPA8 (MA3-014, Thermo Fisher Scientific), anti-PKM2 (#4053, Cell Signaling Technology) and anti-HSP90 (GTX21429, GeneTex, CA). Protein bands were visualized with the Bio-Rad ChemiDOC XRS Imaging system and quantified with ImageJ software.

### 2.4 Lentiviral vectors

pLKO.1-puro lentiviral vectors expressing shRNAs targeting *HSP8A (*sh*HSPA8*_1: NM_006597.3-2040s21c1) *and HSP90AA1* (NM_005348.x-1505s1c1) were purchased from Sigma-Aldrich. These vectors contain a puromycin antibiotic resistance gene for selection of transduced mammalian cells. Full DNA sequence of LAMP-2A was amplified from a pLV-LAMP-2A-GFP plasmid (provided by Ana Maria Cuervo Lab. Primers: Fwd 5’-GGCCCGGGTTGGATCCGCCGCCACCATGGACTACAAAGACGATGACGACAAGGTGTGCTTCCGCC-3’ and Rev 5’-GGAGACGCGTGGATCCTTAAAATTGCTCATA-3’) and cloned into pLV-EF1α-MCS-IRES-Hygro lentiviral vector (Biosettia, CA) to generate LAMP-2A overexpressing construct. Cloning was carried out using the ligation-independent In-Fusion® HD cloning kit (Takara Bio USA, CA). Lentivirus production and transduction were done as described ^33^. Transduced NB4 cell populations were selected with 1.5 µg/mL puromycin for 4 days and knockdown efficiency was assessed by qPCR and western blot analysis.

### 2.5 Granulocytic differentiation assessment

NB4, NB4-R1 and HT93 cells were seeded at a concentration of 0.2×10^6^/mL with 1µM ATRA or dimethyl sulfoxide (DMSO) diluent as the control. Approximately, 1×10^6^ cells were collected on day 2 and 4 to assess CD11b expression and undertake Nitro blue tetrazolium chloride (NBT) assay. Briefly, cells were centrifuged and incubated with PE conjugated anti-CD11b (Dako, Denmark) diluted in FACS buffer (PBS with 1% BSA) for 20 min on ice. DAPI (BioLegend, CA) was used to stain and exclude the dead cells. Signal was acquired by BD FACS LSR II Special Order Research Product (SORP) and analyzed using FlowJo software (BD Biosciences, San Jose, CA). For NBT assay, the cells were incubated in working solution, containing NBT (N5514, Sigma-Aldrich) and phorbol 12-myristate 13-acetate (PMA) (P1585, Sigma-Aldrich) diluted in media for 20 min at 37°C. NBT stained cells were cytospun to the slides and counterstained with Safranin O (HT90432, Sigma-Aldrich). Images were taken using an EVOS microscope (Thermo Fisher Scientific) and NBT stained cells were counted manually.

### 2.6 Photoactivatable CMA reporter assay

NB4 cells were transduced with lentivirus carrying the pSIN-PAmCherry-KFERQ-NE plasmid. pSIN-PAmCherry-KFERQ-NE was a gift from Shu Leong Ho (Addgene plasmid #102365; http://n2t.net/addgene:102365; RRID: Addgene_102365) ^35^. Post-transduction, the cells were selected with 1.5 µg/mL puromycin for 4 days. To activate the mCherry fluorescent protein, cells were exposed to UV light (SOLIS-405C, Thorlabs, NJ, USA) for 7 min, followed by change of media and addition of treatment, where necessary. Cells were collected and stained with LIVE/DEAD™ fixable green (L23101, Thermo Fisher Scientific) to exclude dead cells and fixed with 2% (v/v) paraformaldehyde (PFA), every 24h for three days. mCherry signal was acquired by BD FACS LSR II SORP and analyzed using FlowJo software.

### 2.7 Imaging flow cytometry

Cells were stained with LIVE/DEAD™ fixable green and CD11b-PE as stated before. They were fixed and permeabilized with FOXP3 Fix/Perm Buffer Set (#421401, BioLegend) and stained with anti-LC3B antibody (3868; Cell Signaling Technology). Anti-rabbit Alexa-647 (111-606-045, Jackson ImmunoResearch) was used as secondary antibody. Samples were acquired by the Amnis^®^ ImageStream^®X^ MkII and analyzed with IDEAS™ software (Luminex Corporation, Austin, TX).

### 2.8 Carboxyfluorescein succinimidyl ester (CFSE) assay

Cells were collected, washed and re-suspended in PBS containing 0.1% FBS. CFSE (Sigma-Aldrich) was added to a final concentration of 5µM and cells were incubated at room temperature for 8 minutes, followed by washing three times with PBS containing 10% FBS. The cells were then suspended in media at 0.2 million cells/ml density. Samples were collected at 24h intervals for 3 days and fixed with 2% PFA. CFSE signal was acquired in the FITC channel by BD FACS LSR II SORP and analyzed using FlowJo software.

### 2.9 Vesicle enrichment

Two million NB4 cells treated as mentioned, were collected and washed with PBS. Pellet was re-suspended in 1ml homogenization medium containing protease inhibitor and homogenized using a Dounce homogenizer. The homogenized liquid was collected in Eppendorf tubes and centrifuged at 1000g for 10 minutes at 4°C. The supernatant was transferred to a second tube and centrifuged at 3000g for 10 minutes, followed by final centrifugation step at 17,000g for 15 minutes, after which the supernatant was discarded and the pellet was eluted in homogenization medium containing 4X loading buffer and β-Mercaptoethanol.

### 2.10 Statistical analysis

Nonparametric Mann-Whitney-U tests or two-way ANOVA were applied to compare the difference between two groups using Prism software. P-values <0.05 were considered statistically significant.

## 3 Results

### 3.1 High CMA associated mRNA transcripts are deregulated in AML blasts

To understand the potential dynamic of CMA in AML samples, we analyzed CMA related mRNA transcripts levels in different AML subtypes, healthy HSCs and polymorphonuclear cells (PMN) data from the BloodSpot (Bagger et al. 2016) database. Briefly, during CMA, HSPA8 recognizes client proteins that express the KFERQ-like motif and forms part of the cytosolic chaperone complex composed of HSP90, STIP1 and HIP proteins. Additionally, DNAJ/HSP40 protein and BAG1 regulates the ATPase activity of HSPA8 (Figure 1A). Interestingly, mRNA transcripts encoding proteins of the HSPA8 chaperone complex such as *HSPA8, HSP90AA1, ST13* and *STIP1* are highly expressed in immature hematopoietic stem cells and AML cells compared to mature granulocytes (Figure 1B) suggesting an abundance of CMA players in immature hematopoietic cells. On the other hand, macroautophagy-associated mRNA transcripts such as *MAP1LC3B, WIPI1, WIPI2, GABARAPL1* and *GABARAPL2* are expressed at lower levels in healthy HSCs and AML patients than in polymorphonuclear (PMN) cells (Supplementary Figure 1). This observation supports our earlier report showing higher expression of macroautophagy genes in primary healthy PMN cells than in AML blasts ^37^. These results suggest that the expression of CMA regulating genes are expressed highly in immature cells and downregulated in differentiated healthy myeloid cells.

**Figure 1:**
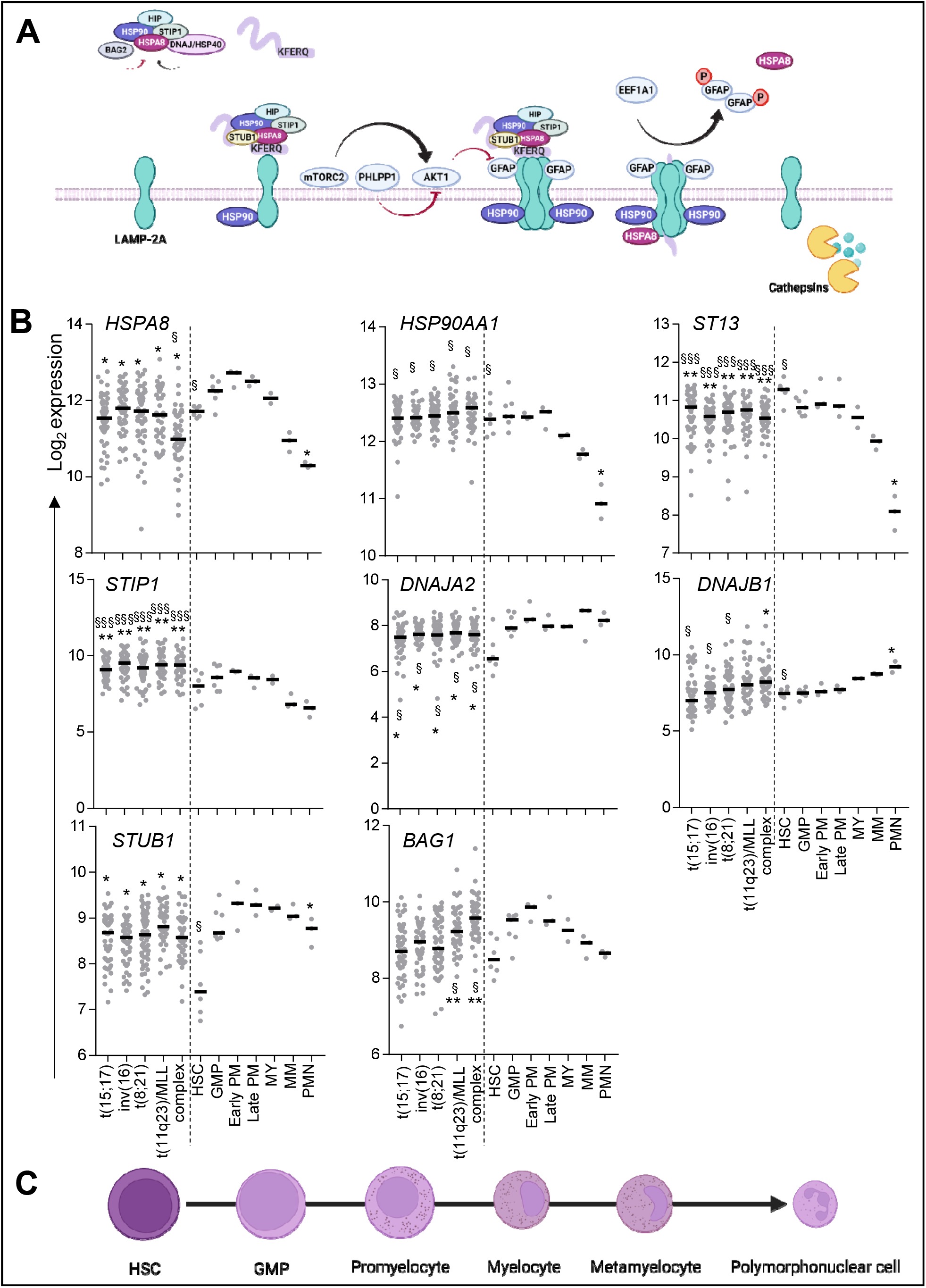
Increased CMA gene expression in AML cells. (**A**) Schematic representation of CMA pathway regulation (BAG2: BAG family molecular chaperone regulator 2; HSP90: Heat shock protein 90; HSP40/DNAJ: Heat shock 40 kDa/DnaJ homologue; STIP1: Stress-induced-phosphoprotein 1; HIP: HSC70-interacting protein; STUB1: E3 ubiquitin-protein ligase CHIP; mTORC2: mammalian target of rapamycin complex 2; PHLPP1: PH domain leucine-rich repeat-containing protein phosphatase 1; AKT1: Protein kinase B; GFAP: Glial fibrillary acidic protein; EF1A1: Elongation factor 1-alpha 1) (**B**). Bloodspot data bank analysis of CMA associated genes’ expression. MWU *=0.05, **p=0.005, ***p<0.001. ns=not significant. n.a=not available.

### 3.2 CMA activity decreases during granulocytic differentiation of healthy and APL cells

We investigated how autophagy gene expression varies in non-malignant myeloid cells undergoing granulocytic differentiation. CD34^+^ cord blood derived human hematopoietic cells were differentiated with GM-CSF for 3 days and G-CSF for up to 7 days. The mRNA levels of CMA-associated splice variant A of *LAMP2* and *HSP90AA1* decreased, whereas mRNA levels of macroautophagy-associated genes *MAP1LC3B* and *WIPI1* increased with time (Figure 2A). Next, we aimed to understand how this phenomenon plays out during leukemic ganulocytic differentiation of AML cells. For this, we used the APL cell lines HT93 and NB4, which can be triggered toward granulocytic differentiation using ATRA. APL cells treated with ATRA for 2 or 4 days showed increased levels of the known CMA client, PKM2, and a decrease in the protein level of the CMA co-chaperone HSP90α (Figure 2B). To confirm that CMA level is reduced upon ATRA therapy, we used LAMP-2A/HASPA8 co-localization ^38^. In line, we observed significantly less co-localization between HSPA8 and LAMP-2A after ATRA treatment (Figure 2C). To validate that these observations indicate CMA activity, we used a CMA reporter assay ^35,39^. Briefly, NB4 APL cells were transduced with a plasmid containing a photoactivatable mCherry tagged KFERQ motif sequence. Once mCherry is activated, degradation of the mCherry-KFERQ CMA client can be monitored by measuring mCherry intensity via dots formation or flow cytometry (Figure 2D). Unfortunately, AML blasts contain very little cytoplasm. This results in low-resolution images of mCherry-KFERQ dots, which are impossible to quantify (Supplementary Figure 2A). Another issue is that they are suspension cells and do not grow in a monolayer. This causes not all cells receive UV uniformly to activate mCherry. We therefore used mCherry fluorescence intensity as a way to measure KFERQ degradation using flow cytometry. This also allowed us to measure degradation over a longer period. We collected samples every 24h for 3 days and observed that ATRA-treated samples had a lower rate of degradation compared to DMSO treated samples. mCherry signal accumulation upon Bafilomycin A1 treatment confirmed that this degradation is mediated by lysosomes as Bafilymycin A1 is a V-ATPase inhibitor which inhibits lysosomal degradation by keeping the luminal pH high (Figure 2E). In addition, we observed accumulation of ATG5-ATG12 complex and DAPK2 with ATRA treatment along with increased LC3B flux, which is in line with our previous findings that demonstrated increase in macroautophagy flux upon ATRA treatment (Supplementary Figure 2B-C) ^37,40^. Interestingly, knocking down *ATG5*, which reduces macroautophagy flux as well as the rate of differentiation in APL cells ^37^ abolishes the drop in CMA caused by ATRA treatment (Supplementary Figure 2D-E).

**Figure 2:**
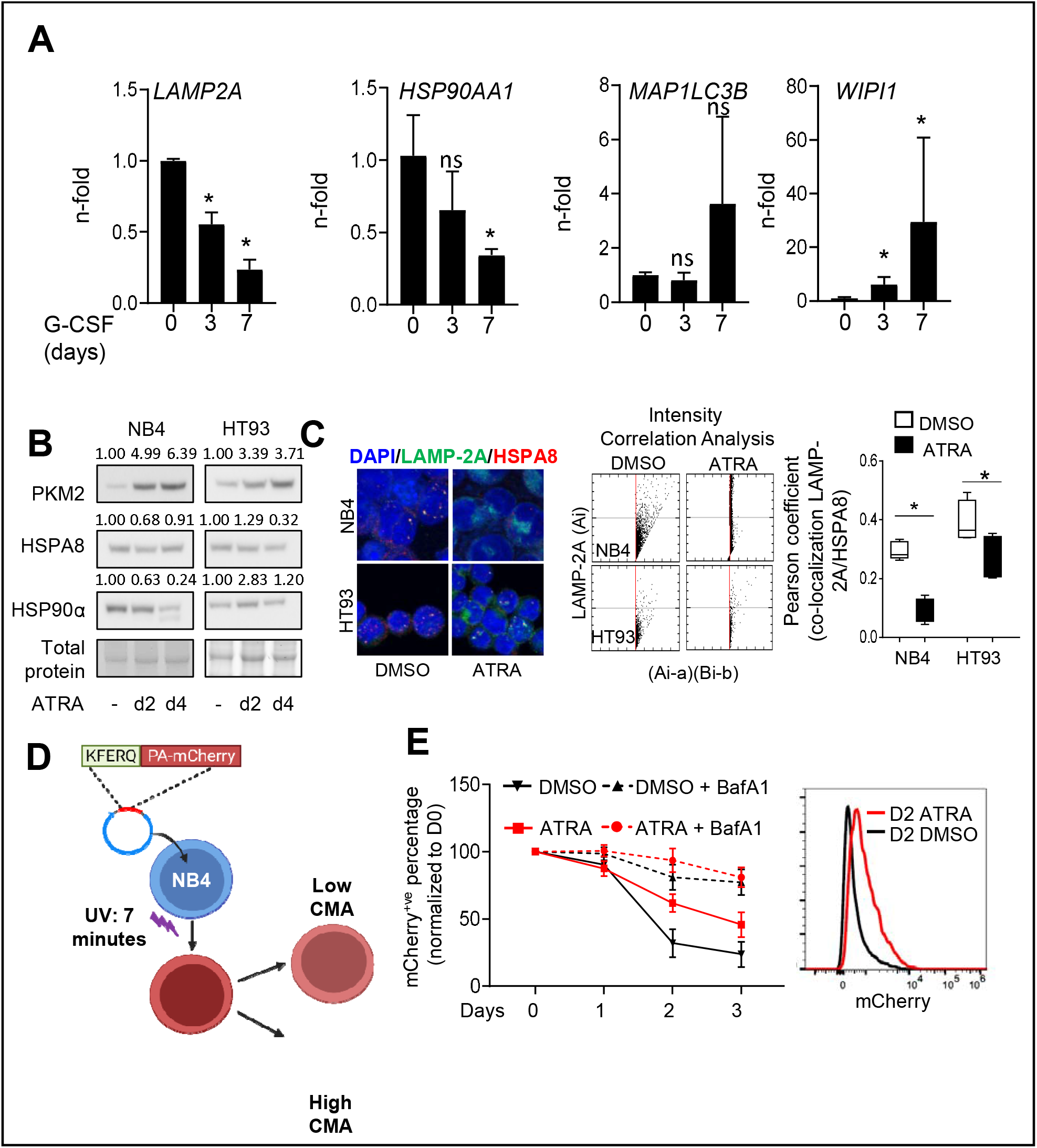
CMA related gene expression and CMA activity decreases during granulocytic differentiation. (**A**) Human CD34^+^ hematopoietic cells were differentiated towards granulocytes using GM-CSF and G-CSF at the indicated time point. Transcript levels of CMA related gene *LAMP2* (variant A) and *HSP90AA1* and macroautophagy related *MAP1LC3B* and *WIPI1* were analyzed by qPCR. Values were normalized to the *HMBS* housekeeping gene. Results of at least two independent experiments are shown as n-fold change compared with day 0 treated cells. (**B**) APL cell lines NB4 and HT93 were treated with ATRA (1µM) for 2 and 4 days. A. Protein extracted from whole cell lysate was subjected to immunoblotting using anti-HSPA8, anti-HSP90 and anti-PKM2 antibodies. Total protein is shown as a loading control (N=3) **(C)** NB4 and HT93 cells were treated with ATRA for 2 days and fixed and stained for endogenous LAMP2A and HSPA8. Degree of co-localization was analyzed using the JaCoP Plugin on ImageJ software. (N=3) (**D**) Scheme showing the experimental design to interpret CMA activity in cultured cells. (**E**) NB4 cells were transduced with lentivirus carrying the pSIN-PAmCherry-KFERQ-NE plasmid and treated with ATRA (1uM) with or without BaA1 (50nM). mCherry intensity was measured by flow cytometry. Left: percentage of mCherry positive cells. D0 was set at 100%. Values are the mean of four biological replicates. Right: histogram of mCherry intensity upon ATRA treatment (red) or in DMSO treated cells (black) on day 2 created on FlowJo.

Taken together, our data prove that high expression of CMA-associated genes accounts for high CMA activity, which decreases upon differentiation of myeloid cells towards neutrophils.

### 3.3 Macroautophagy activation primes the cells to differentiate

While macroautophagy has been frequently linked to ATRA-induced differentiation, of APL cells, it is still unclear if macroautophagy is the cause or the consequence of differentiation. Therefore, we followed the kinetics of macroautophagy activity during granulocytic differentiation, using ImageStream^®^ imaging flow cytometer, which integrates the features of flow cytometry and fluorescence microscopy combined with advanced image analysis methodology. This allowed a more precise assessment of macroautophagy of myeloid cells at different stages of differentiation. NB4 cells were treated with ATRA for 2 days and co-stained for LC3B and the myeloid differentiation marker, CD11b. We analyzed the simultaneous expression of CD11b and LC3B in cells, using *IDEAS*™ software. Dead cells were excluded using LIVE/DEAD fixable green (FITC^+^) and only cells in the focus stream were considered for analysis (Supplementary Figure 3A-D). The mask threshold was determined based on a single cell out of CD11b^+^ population and applied on the rest of the samples. We counted the number of LC3B dots in presence and absence of Bafilomycin A1 and calculated macroautophagy flux by subtracting the number of dots in untreated samples from the treated ones. Using this protocol, we were able to detect an increased macroautophagy flux (BafA1^+^-BafA1^-^) in ATRA treated cells compared to DMSO, validating the robustness of the method. Interestingly, increased macroautophagy flux was already detected prior to CD11b expression. The autophagy flux increased steadily, but not significantly, upon further myeloid maturation of the cells according to their CD11b expression levels (low, medium or high) (Figure 3A). This suggested that AML cells already experienced increased macroautophagy flux when primed for differentiation (ATRA treated CD11b negative cells). To test our hypothesis, we utilized the ATRA-resistant NB4-R1 cells. NB4-R1 cells are unique because they respond to retinoic acid transcriptionally, upregulate RARA, down-regulate RXR, degrade PML-RARA, and become competent for maturation (Duprez, Lillehaug, Naoe, et al. 1996b; Duprez, Lillehaug, Gaub, et al. 1996; Duprez, Lillehaug, Naoe, et al. 1996a; Pendino et al. 2001) but only fully differentiate upon addition of cyclic adenosine monophosphate (cAMP). We treated NB4 cells for 48h with ATRA and ATRA + cAMP and measured macroautophagy flux. It was observed that ATRA treatment alone activated macroautophagy in NB4-R1 cells without inducing differentiation (Figure 3B) while cAMP did not have an additive effect. These data strongly suggest that macroautophagy is upregulated to prime APL cells towards granulocytic differentiation but cannot lead to full maturation. We already observed that CMA activity decreases during differentiation. Therefore, we hypothesized that CMA could be the driving factor trigger maturation of these cells. We assessed LAMP-2A and HSPA8 co-localization in NB4-R1 cells upon combined ATRA and cAMP treatment. The degree of co-localization was significantly reduced when NB4-R1 cells were terminally differentiated with the addition of cAMP (Figure 3C). Overall, these observations with NB4-R1 cells showed that macroautophagy activation precedes differentiation and primes APL cells, whereas CMA attenuation happens at a later stage to drive final maturation (Figure 3D).

**Figure 3:**
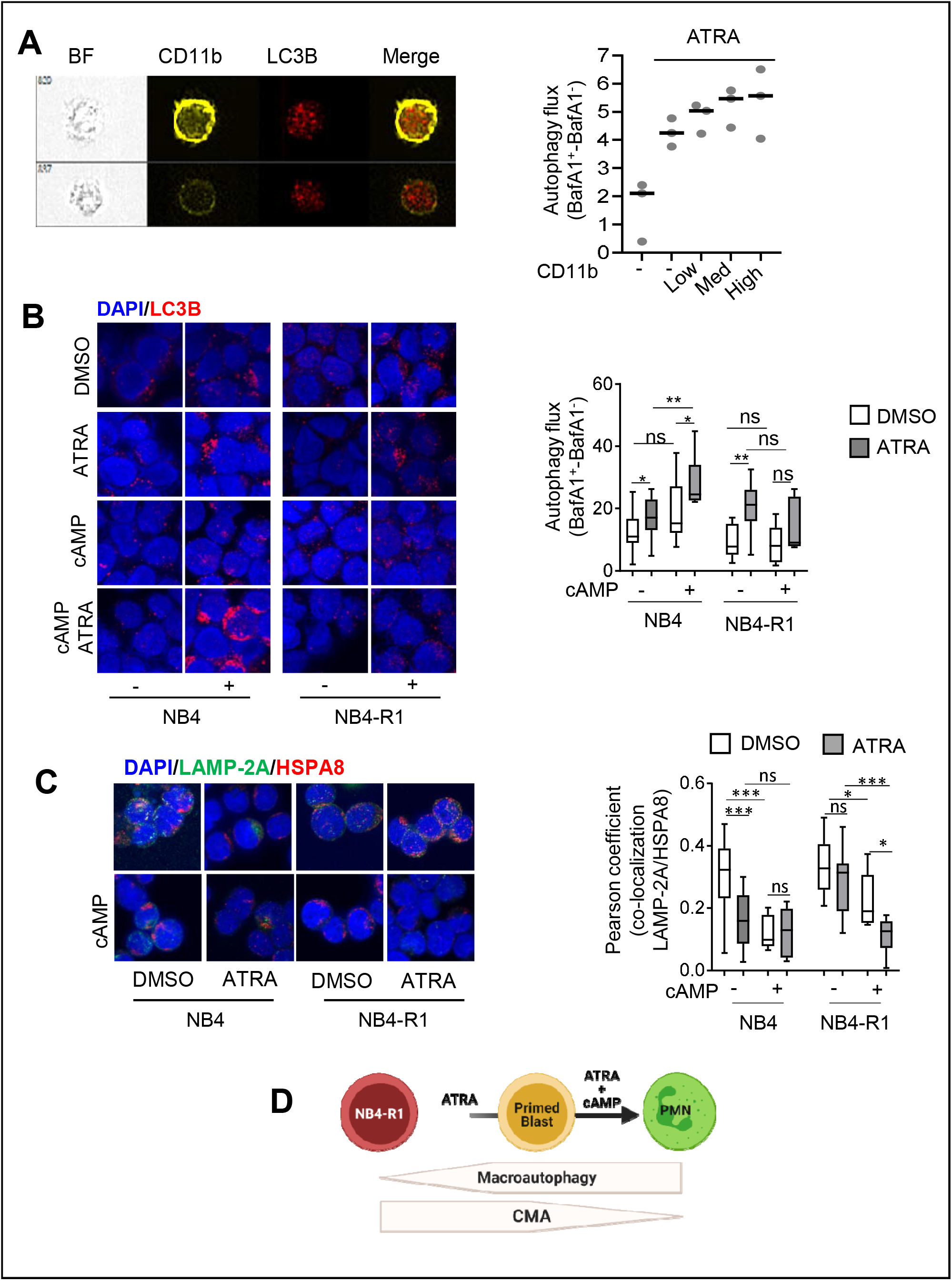
Macroautophagy primes APL cells for differentiation. (**A**) Left: Snapshot of imaging flow cytometry analysis using our protocol to combine surface staining (CD11b) and intracellular staining (LC3B). NB4 cells were treated with ATRA to induce differentiation and subjected to surface staining CD11b-PE together with intracellular staining LC3B. Images were acquired using the 40x objective lens. BF: Bright Field. Right: Quantification of the macroautophagy flux (BafA1^+^-BafA1^-^) upon ATRA treatment classified based on CD11b expression level. (**B**) NB4-R1 Cells treated with ATRA in presence or absence of cAMP (100µM) were stained for endogenous LC3B and images were acquired by confocal microscope. LC3B dots were quantified using ImageJ software and the flux was calculated (BafA1^+^-BafA1^-^). **(C)** NB4-R1 Cells treated with ATRA with or without cAMP for 2 days were subjected to endogenous LAMP2A and HSPA8 immunofluorescence analysis. Pearson’s Coefficient was used to determine degree of co-localization (N=3). (D) Scheme demonstrating the findings of the experiments with NB4-R1 cells. Macroautophagy activity primes APL blasts towards maturation, followed by CMA downregulation to lead to neutrophils.

### 3.4 LAMP-2A ectopic expression decelerate APL differentiation

With the aim of understanding the change in CMA during granulocytic differentiation better, we wondered if modulating CMA in NB4 cells affects granulocytic differentiation. To quantify granulocytic differentiation we measured CD11b surface marker expression by flow cytometry and analyzed the percentage of CD11b^+^ cells using the FlowJo software as an indication of increased or decreased rate of differentiation. CD11b is expressed during myeloid differentiation and gradually increases upon differentiation. As a functional assay, we used nitroblue tetrazolium (NBT). The oxidative respiratory burst in neutrophils produce reactive oxygen species (ROS) which can reduce the yellow NBT into blue formazan deposits that can be quantified from microscopy images and serves as an indication of complete maturation ^45^. LAMP-2A has been described in most literature as the limiting factor of CMA in other models. Therefore, we started by ectopically expressing LAMP-2A in NB4 cells (Figure 4A). We confirmed a significantly higher CMA activity in LAMP-2A overexpressing NB4s compared to control cells within 24 hours by measuring KFERQ degradation (Figure 4B). To confirm that the mCherry reduction was not due to aberrant proliferation of high LAMP2A expression, we also measured proliferation via Carboxyfluorescein succinimidyl ester (CFSE) assay (Supplementary Figure 4A). LAMP-2A overexpressing cells did not show any difference in their rate of proliferation, which indicates that the change in mCherry signal did not arise from a difference in rate of cell division. In line, LAMP-2A overexpressing cells showed a lowered rate of differentiation upon ATRA treatment compared to the control cells using two independent assays (Figure 4C, D) suggesting that high CMA activity impedes ATRA-induced differentiation.

**Figure 4:**
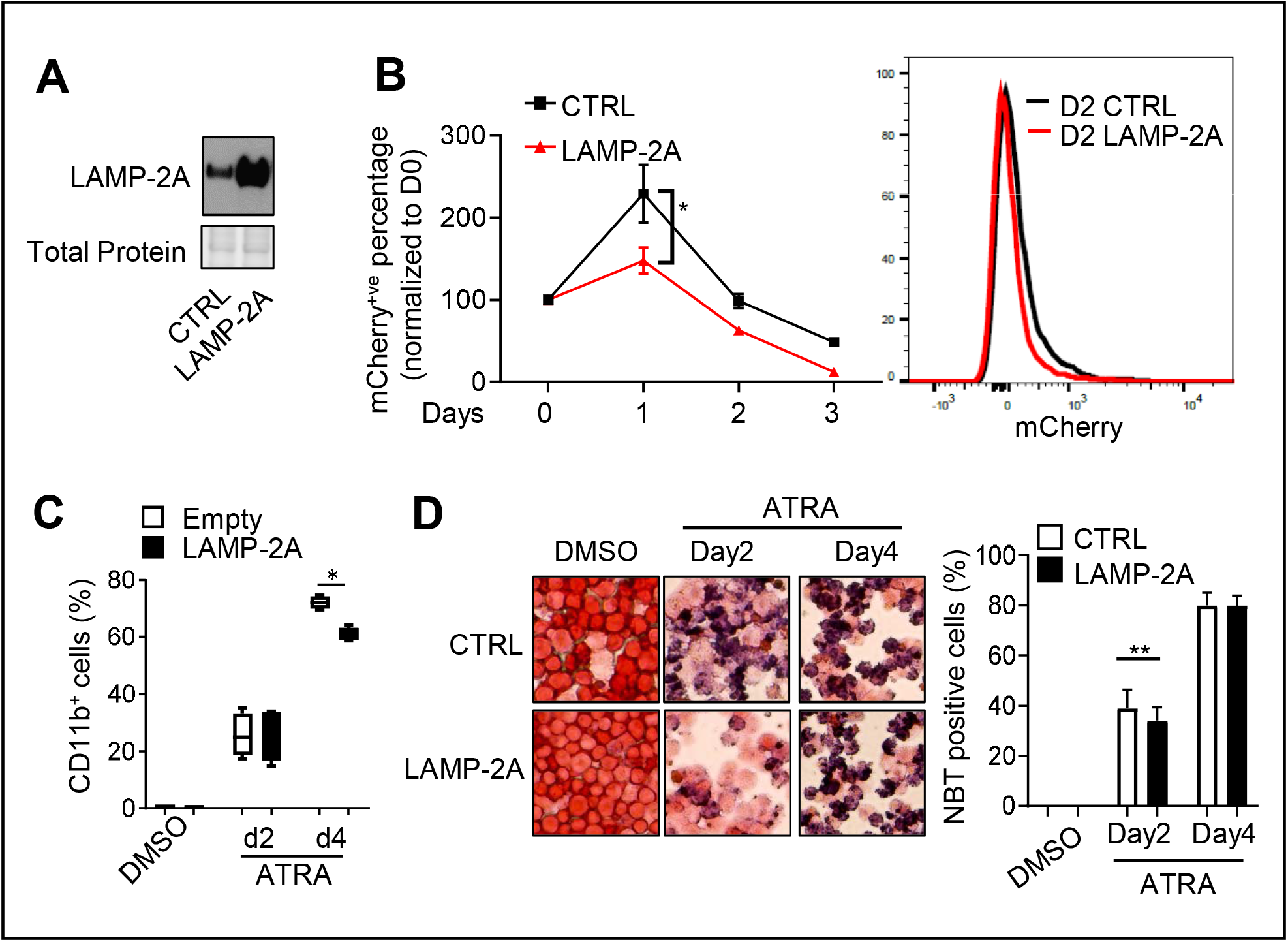
Increased CMA activity attenuates APL differentiation. (**A-D**) LAMP2A was ectopically overexpressed in NB4 cells. (**A**) Overexpression of LAMP2A confirmed by western blot. (**B**) CMA activity was analyzed using the PA-mCherry-KFERQ reporter as described in figure 3D (N=2). (**C**) LAMP2A overexpressing and control NB4 cells were treated with ATRA for 2 and 4 days. Granulocytic differentiation was determined with CD11b surface marker expression by flow cytometry and **(D)** NBT assay (N=3).

Altogether, we found that increased LAMP-2A leads to high CMA activity, which slows down ATRA-induced differentiation. However, even though the same phenotype is observed in a LAMP-2A depleted model, it cannot be attributed to CMA, as the activity remains unaffected.

### 3.5 HSPA8 is the determinant factor for CMA activity in APL cells

Next, we focused on *HSPA8 and HSP90AA1*, two different key CMA regulators, and analyzed if their depletion (Figure 5A, E) had any effect on CMA. We found that knocking down *HSPA8* reduced CMA activity (Figure 5B). *HSPA8* knockdown cells showed no significant change in CD11b marker expression upon ATRA treatment (Figure 5C). Importantly, NBT reduction assays, which detect oxidative bursts in functional neutrophils, showed that ATRA treatment of cells with lower CMA activity resulted in more mature granulocytes (Figure 5D). In parallel, knockdown of *HSP90AA1* showed decreased co-localization of HSPA8 and LAMP-2A and accelerated differentiation based on CD11b expression (Figures 5F,G).

**Figure 5:**
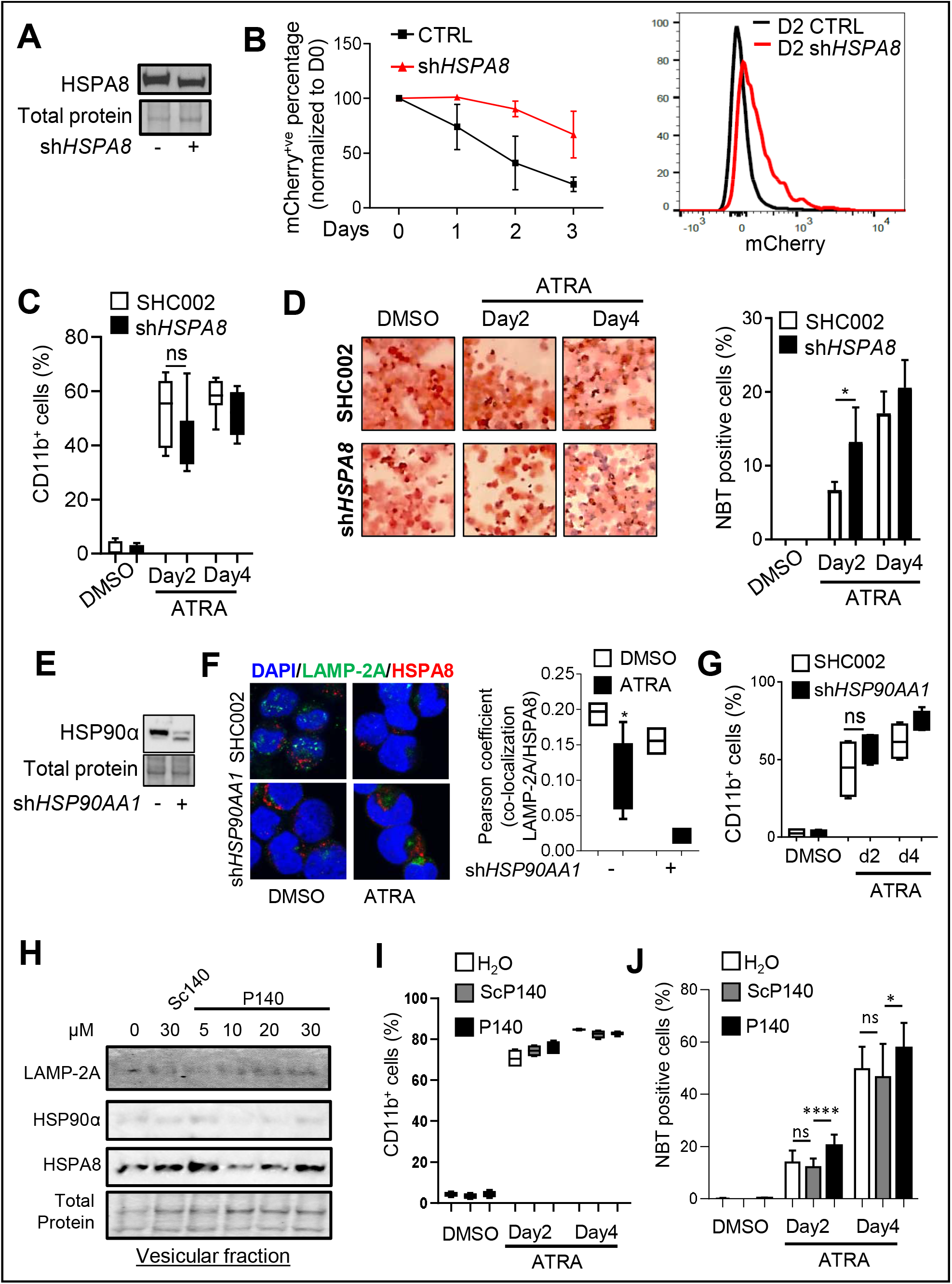
Lowering CMA activity supports ATRA-mediated APL differentiation. (**A-D**) NB4 cells were stably transduced with non-targeting shRNA (SHC002) or shRNAs targeting *HSPA8* lentiviral vectors and differentiated with 1μM ATRA for 2 or 4 days. (**A**) HSPA8 western blot analysis of control and sh*HSPA8* expressing NB4 cell populations. (**B**) CMA activity in NB4 control (SHC002) and *HSPA8* knockdown cells (N=2). (**C**) Flow cytometry analysis of CD11b surface expression NB4 control (SHC002) and *HSPA8* knockdown NB4 cells upon ATRA treatment. Values are the mean of at least two biological replicates. (**D**) NBT reduction in ATRA-treated NB4 control (SHC002) and *HSPA8* knockdown cells. Left panel: Representative images of NBT assays in control and HSPA8 depleted NB4 cells. Right panel: Quantification of the percentage of NBT^+^ cells (N=3) (**E-G**) NB4 cells were stably transduced with non-targeting shRNA (SHC002) or shRNAs targeting *HSP90AA1* lentiviral vectors and differentiated with 1μM ATRA for 2 or 4 days. (**E**) Western blot analysis to conform HSP90 knockdown. (**F**) LAMP2A-HSPA8 co-localization was captured by immunofluorescence microscopy and quantified using JaCoP Plugin on ImageJ software. (N=3) (**G**) Flow cytometry analysis of CD11b surface marker expression NB4 control (SHC002) and *HSP90AA1* knockdown NB4 cells upon ATRA treatment **(H)** NB4 cells were treated with different concentration of P140 for 12h and subjected to vesicular enrichment protocol. Enriched vesicles were analyzed for CMA markers LAMP2A, HSP90α and HSPA8 by western blot. (**I-J**) NB4 cells were co-treated with 1µM ATRA together with P140 (10µM), ScP140 (10µM) or water. (**I**) Flow cytometry analysis of CD11b surface marker expression. (**J**) NBT reduction in ATRA-treated NB4 cells (N=3).

In addition to genetically inhibiting CMA, we used P140/Lupuzor™, a peptide drug that is currently in clinical trial for treatment against systemic lupus erythematosus (SLE), as a way to inhibit CMA pharmacologically. P140 binds to HSPA8 and destabilizes the LAMP-2A-co-chaperone complex on the lysosomal membrane ^46–48^. We treated NB4 cells with different concentrations of P140 and determined the amount of LAMP-2A, HSPA8 and HSP90α on the enriched vesicular fraction of the cells. We determined 10µM to be the optimum concentration to disrupt the co-chaperone complex. The scrambled peptide (ScP140) used as control did not have any effect on the co-chaperone proteins (Figure 5H). Combined treatment of P140 and ATRA did not affect CD11b expression but we observed accelerated terminal differentiation using NBT assay, similar to our results with the *HSPA8* knockdown cells (Supplementary Figure 5B; Figure 6I-J).

Overall, our data indicate that HSPA8 and the accompanying co-chaperones are limiting factors in CMA regulation during ATRA mediated APL differentiation and even minor changes in their expression levels affect granulocytic differentiation in NB4 APL cells.

## 4 Discussion

The importance of autophagy in hematopoietic stem cell maintenance has been demonstrated in earlier studies ^9,31^. Both macroautophagy and CMA contribute to preserving healthy HSCs by regulating metabolic pathways. Selective macroautophagy processes such as, mitophagy and lipophagy, bring about the shift in energy metabolism needed for myeloid differentiation ^49^ and CMA specifically targets the polyunsaturated fatty acid metabolism pathway to regulate activation of HSCs ^31^. Leukemic cells, by nature, have blocked differentiation and only the AML subtype, APL responds efficiently to differentiation therapy. Studies have shown the importance of macroautophagy in this type of differentiation but so far, there has been no report of how CMA might influence this process. Here, we have established that expression of CMA related genes lowers during normal and APL differentiation and the effectivity of ATRA is facilitated by a decrease in CMA activity. Additionally, we have demonstrated the importance of the HSPA8 chaperone complex in this process and recognized its role as a limiting factor in CMA regulation during APL differentiation.

Owing to its connection with ageing, CMA has been studied more extensively in the pathophysiology of neurodegenerative diseases. However, research in the past decade have pointed towards its relevance in cancer as well. CMA markers are rather highly expressed in various solid cancer types, and CMA activity is frequently upregulated in cancer cells compared to normal cells ^27,50,51^. Interestingly, in most cases, high CMA activity is observed regardless of the macroautophagy status of the cells. In the present work, we report that acute myeloid leukemia cells also exhibit high CMA in terms of both gene expression as well as activity. As APL cells mature, CMA activity decreases contrasting the increase of macroautophagy during the same process. It is unclear whether the two processes are co-regulated or whether this reciprocal phenotype is observed independently. Of interest, CMA is under the control of mechanistic target of rapamycin (mTOR)C2 whereas macroautophagy is mediated by mTORC1 inhibition. Even though these are two separated complexes, the protein kinase B or Akt1 is involved in both mechanisms ^52^. Therefore, the possibility exists that a common player upstream of both pathways regulates them simultaneously.

Wide ranged CMA-related studies in multiple model systems have been limited in parts, due to the difficulty in quantifying the process. There are far fewer assays to measure CMA activity directly and most of them rely on indirect readouts such as, CMA related genes expression or protein level of known CMA clients. The only specific way to measure CMA activity is by using a fluorescent reporter tagged to KFERQ motif. It also serves as a way to differentiate CMA from endosomal microautophagy (eMI) because this reporter is only targeted to CMA competent lysosomes and not endosomes ^39,53^. Unfortunately, the requirement of activation with UV light makes *in vitro* studies rather complicated due to the UV sensitivity of cells. We were able to optimize this assay in our leukemia cell lines and showed that even a small decrease in HSPA8 levels is sufficient to affect CMA activity in NB4 cells, which results in accelerated maturation of APL blasts upon ATRA treatment. HSPA8 has been observed to be involved in certain hematological malignancies. For example, HSPA8 expression contributes to the leukemic phenotype in chronic myeloid leukemia ^54^. Proteomic analysis identified HSPA8 to be differentially expressed in HSC-like cell fraction isolated from patients with hematological malignancies ^55^. Nevertheless, no previous report has shown its connection to CMA modulation during APL differentiation.

Roughly, half of all proteins contain at least one canonical KFERQ-like motif ^56^ but not all KFERQ motif containing proteins are degraded by CMA. Harboring the motif only denotes the possibility of being degraded via CMA. Only a handful of these proteins have been experimentally validated. Among these, PKM2, HK2 and AF1Q could be prospective candidates to affect myeloid differentiation ^32^. Since differentiation is a tightly regulated process, it will be of interest to explore client candidates of macroautophagy and CMA whose degradation could contribute to ATRA-mediated differentiation. This is not a very straightforward process because change in CMA activity affects the metabolic landscape of cancer cells, which favors degradation of certain clients over others to allow the cells to sustain Warburg effect ^28^. Therefore, it is difficult to pinpoint if the phenotype is a result of degradation of a specific client or the change in cellular processes that resulted from CMA activity perturbation.

ATRA treatment in combination with chemotherapy or ATO has high success rate in APL treatment, but resistance to this treatment can occur. Various mechanisms of resistance development have been elucidated including mutations in the PML/RARα ligand binding domain and RARE domain, epigenetic regulation of RARα promoter, high expression of MN1 transcription factor and release of retinoic acid degrading factors by the BM microenvironment ^57,58^. Our findings suggest HSPA8 driven CMA as an inhibitory pathway in ATRA-induced differentiation. With the development of specific CMA inhibitors such as P140/Lupuzor™, it might be possible to target CMA in a clinical setting to increase the effectiveness of ATRA treatment.

## 6 Author Contributions

SR and MH wrote the manuscript. SR, IM, and MH designed and performed experiments, analyzed and interpreted data. NJN and BET provided CD34+ cells and critical input. StM provided critical input and support for the ImageStream® protocol and analysis. AJ, GR and PA provided critical input. SM provided the P140 and ScP140 peptides and critical input. MH and MPT obtained funding, supervised the project and data analysis, and gave the final approval for the publication. All authors discussed the results and commented on the manuscript.

## 8 Acknowledgments

Deborah Shan-Krauer is gratefully acknowledged for excellent technical support. SM would like to acknowledge the French Centre National de la Recherche Scientifique, Région Grand-Est, the Strasbourg University program ITI 2021-2028 (Idex ANR-10-IDEX-0002/STRAT’US project ANR-20-SFRI-0012) and the University of Strasbourg Institute for Advanced Study (USIAS), as well as the Club francophone de l’autophagie (CFATG). All schematic representations were created with Biorender.com.

