## Supplementary Figure Legends for "HSPA8 chaperone complex drives chaperone-mediated autophagy regulation in acute promyelocytic leukemia differentiation"

**Supplementary Figure 1. Related to Figure 1**

Bloodspot data bank analysis of macroautophagy associated genes’ expression: *MAP1LC3B, WIPI1, WIPI2, GABARAPL1, GABARAPL2*. HSC: Hematopoietic stem cell, GMP: Granulocyte monocyte progenitors, early_PM: Early Promyelocyte, late_PM: Late Promyelocyte, MY: Myelocyte, MM: Metamyelocytes, PMN: Polymorphonuclear cells MWU ns=not significant. n.a=not available compared to HSC *p<=0.05, **p<=0.005, ***p<0.001. MWU compared to PMN §p<0.05, §§p<0.005, §§§p<0.001

**Supplementary Figure 2. Related to Figure 2**

(**A**) Confocal images of NB4 cells containing photoactivatable mCherry-KFERQ construct. mCherry was activated using UV light and images were acquired 5 hours after activation (**B-C**) Macroautophagy upregulation upon ATRA treatment in APL cell lines NB4 and HT93. (**A**) Cells were treated with 1µM ATRA for 2 and 4 days and subjected to Western blot analysis of ATG12-ATG5 complex and DAPK2. (**B**) Cells treated with ATRA for 2 days with or without Bafilomycin A1 (200nM, 2h) were immunostained for LC3Band analyzed for macroautophagy flux (BafA1^+^+BafA1^-^). Results shown are average of at least two biological replicates. (**C-D**) Reducing macroautophagy flux keeps CMA level high upon ATRA treatment. (**C**) NB4 cells were stably knocked down with lentivirus containing shRNA targeting *ATG5*. These cells were then treated with ATRA (1uM) for 2 or 4 days and subjected to Western blot analysis of LAMP2A, PKM2, ATG12-ATG5, and DAPK2. Total protein has been shown as loading control. (**D**) Control cells and *ATG5* knockdown cells were stained for HSPA8 (red) and LAMP2A (green). Pearson’s coefficient was quantified to measure CMA (N=3).

**Supplementary Figure 3. Related to Figure 3**

(**A-D**) Gating strategy for ImageStream^®^ data analysis on IDEAS™ software.

**Supplementary Figure 4. Related to figures 4 and 5**

(**A-B**) CFSE staining was used to measure proliferation rate of NB4 cells with overexpression of LAMP2A (A), knockdown of HSPA8 (B) (**C-D**) HSPA8 knockdown and LAMP2A overexpression in HT93 cells. (**C**) Western blots confirming HSPA8 knockdown and LAMP2A overexpression in HT93 cells. (**D**) Flow cytometry analysis of CD11b surface marker in HT93 cells harboring sh*HSPA8* and LAMP2A upon 2 and 4 days of ATRA treatment. (**E**) NBT reduction in LAMP2A overexpressing HT93 cells upon 2 and 4 days of ATRA treatment. Results shown are mean of three biological replicates.

**Supplementary Figure 5. Related to figure 5**

(**A**) Images showing LC3B dots used for macroautophagy flux analysis in sh*HSP90AA1* harboring NB4 cells and subsequent quantification. (**B**) Images showing NBT reduction in NB4 cells treated with ATRA and P140, ScP140 or water for 2 and 4 days.
