## Supplementary Figures for "HSPA8 chaperone complex drives chaperone-mediated autophagy regulation in acute promyelocytic leukemia differentiation"

### Slide 1
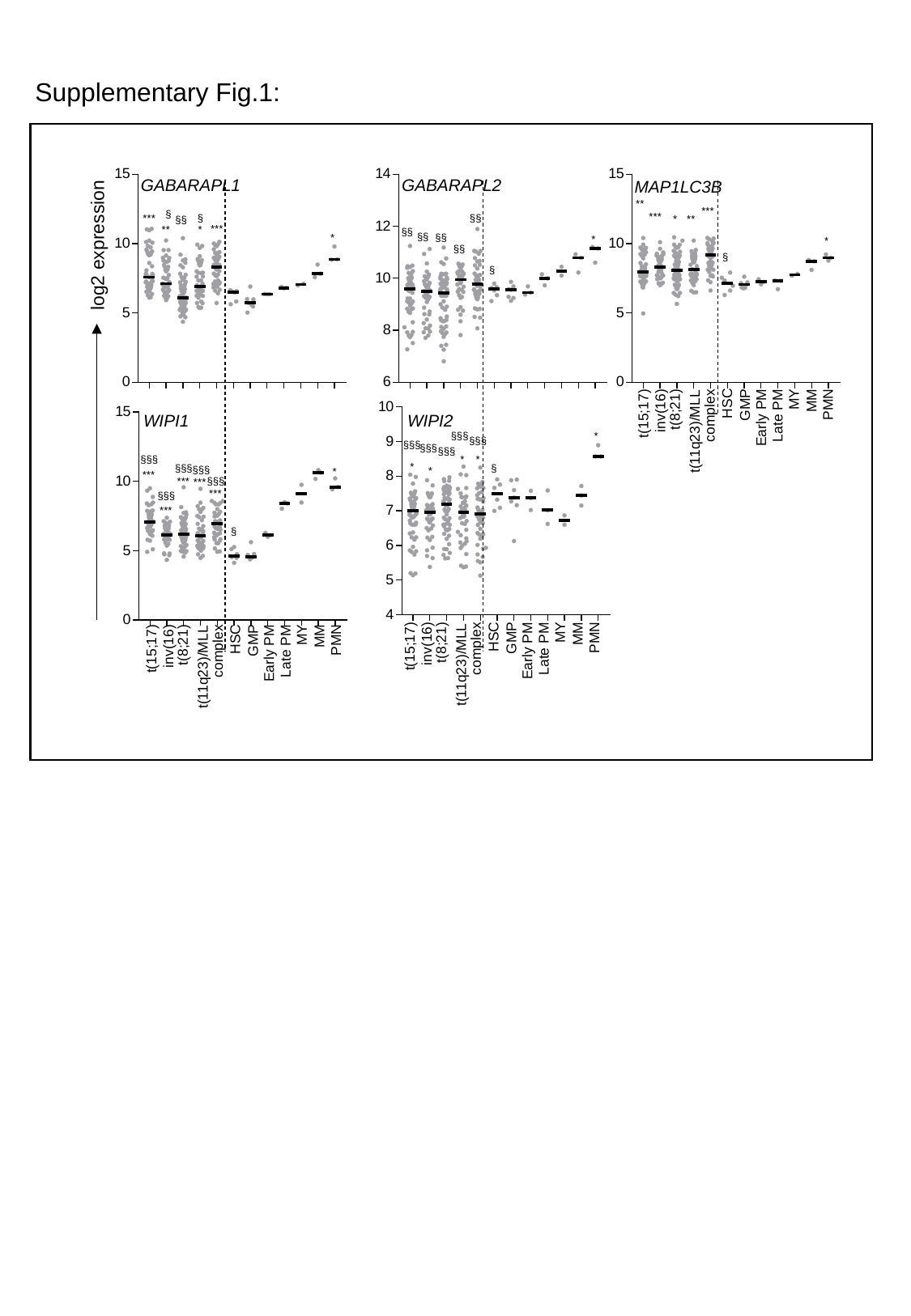

Supplementary Fig.1:
GABARAPL1
GABARAPL2
MAP1LC3B
**
***
§
***
***
§§
§
**
*
§§
***
**
*
§§
§§
§§
*
*
log2 expression
*
§§
§
§
MY
MM
HSC
PMN
GMP
t(8;21)
inv(16)
t(15;17)
complex
Late PM
Early PM
WIPI1
WIPI2
t(11q23)/MLL
§§§
*
§§§
§§§
§§§
§§§
§§§
*
*
*
§
§§§
§§§
*
*
***
***
§§§
***
***
§§§
***
§
MY
MM
MY
MM
HSC
PMN
GMP
HSC
PMN
GMP
t(8;21)
inv(16)
t(8;21)
inv(16)
t(15;17)
complex
Late PM
t(15;17)
Early PM
complex
Late PM
Early PM
t(11q23)/MLL
t(11q23)/MLL

### Slide 2
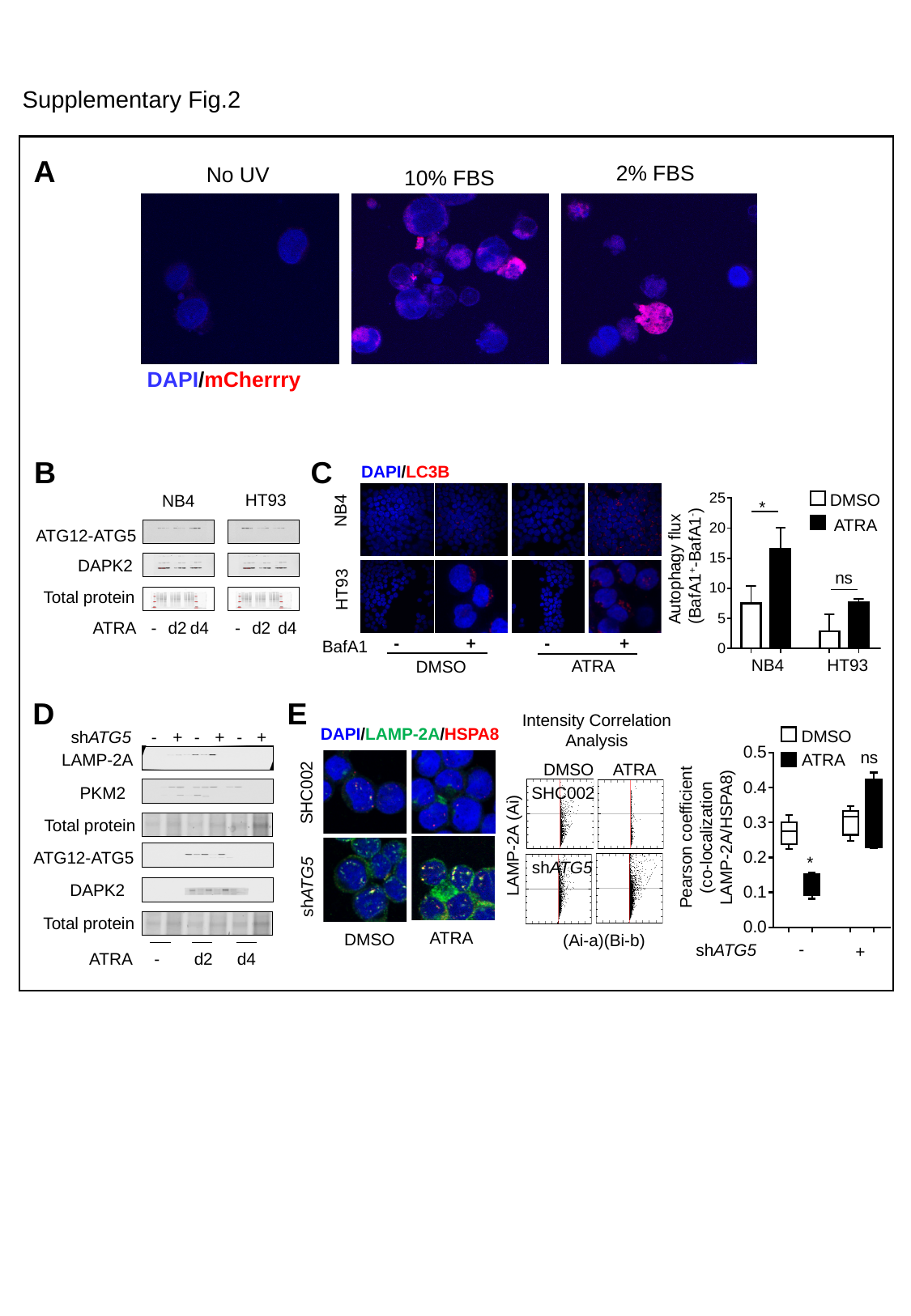

Supplementary Fig.2
A
2% FBS
No UV
10% FBS
DAPI/mCherrry
B
C
DAPI/LC3B
DMSO
HT93
NB4
*
NB4
ATRA
ATG12-ATG5
Autophagy flux
(BafA1^+^-BafA1^-^)
DAPK2
ns
HT93
Total protein
ATRA
-
d2
d4
-
d2
d4
-
+
-
+
BafA1
HT93
NB4
ATRA
DMSO
E
D
Intensity Correlation Analysis
ATRA
DMSO
SHC002
LAMP-2A (Ai)
shATG5
(Ai-a)(Bi-b)
DAPI/LAMP-2A/HSPA8
DMSO
shATG5
-
+
-
+
-
+
ns
LAMP-2A
ATRA
SHC002
PKM2
Pearson coefficient
(co-localization LAMP-2A/HSPA8)
Total protein
ATG12-ATG5
*
shATG5
DAPK2
Total protein
ATRA
DMSO
-
shATG5
+
ATRA
-
d2
d4

### Slide 3
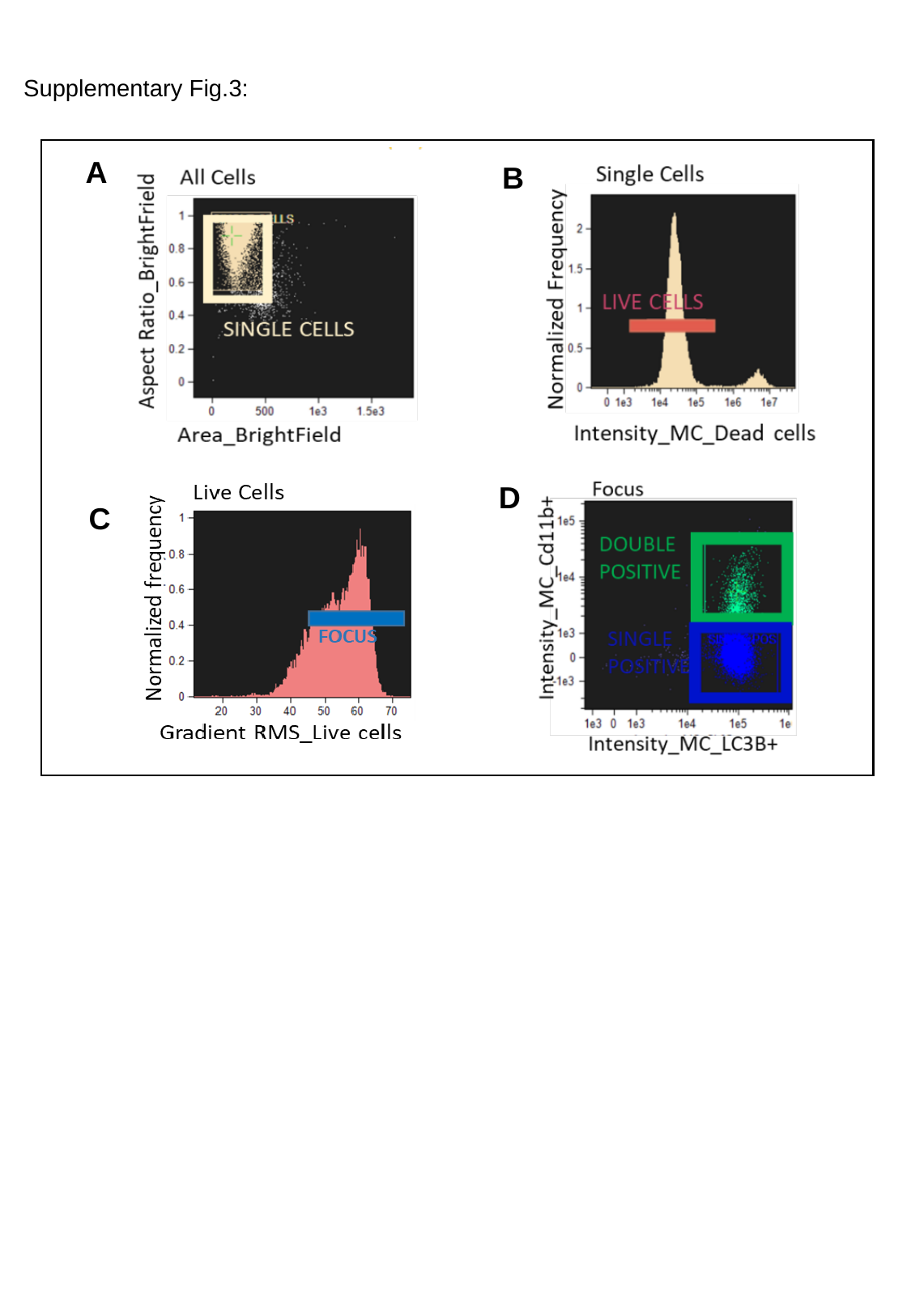

Supplementary Fig.3:
A
B
D
C

### Slide 4
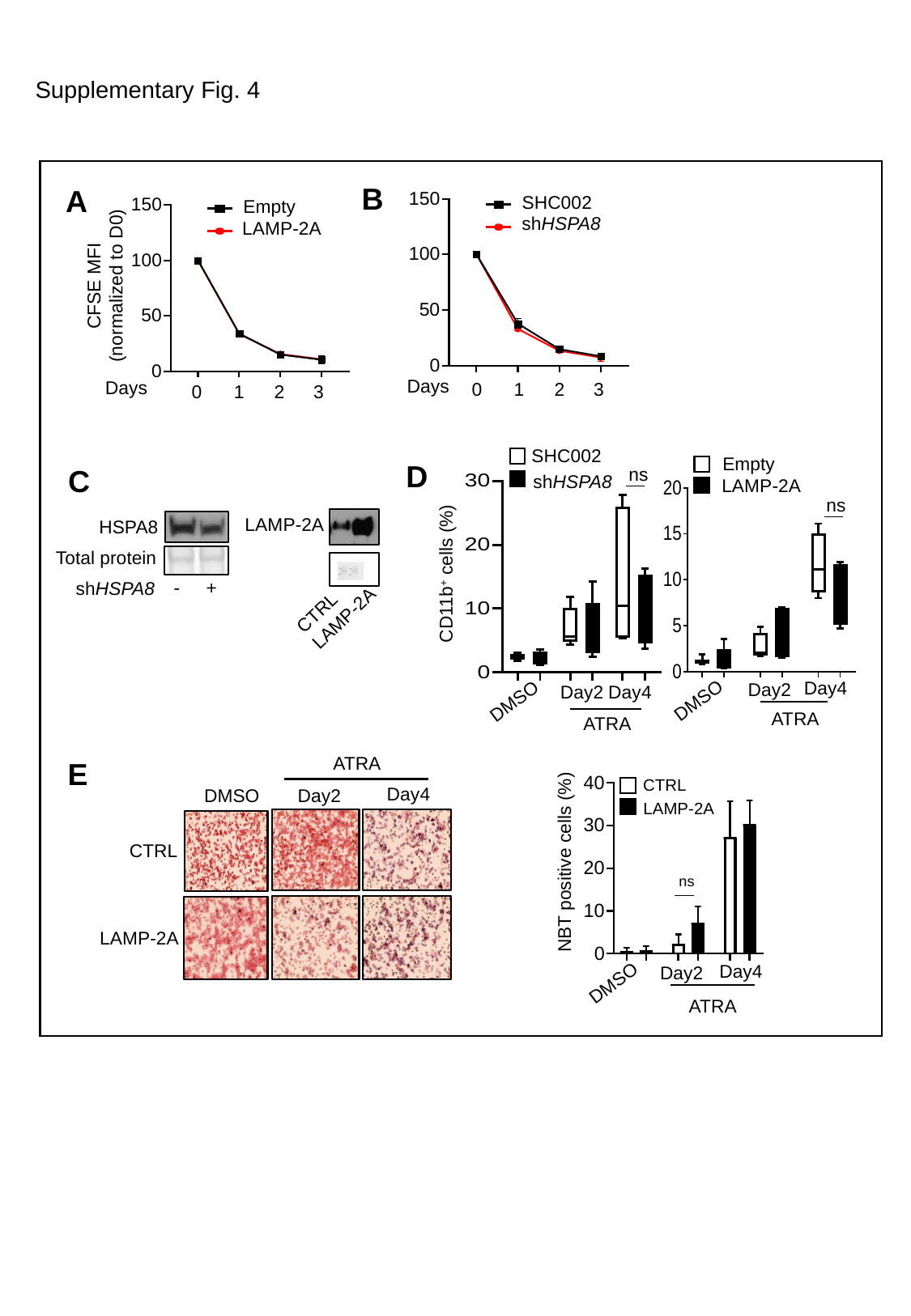

Supplementary Fig. 4
B
A
SHC002
Empty
shHSPA8
LAMP-2A
CFSE MFI
(normalized to D0)
Days
Days
0
1
2
3
0
1
2
3
SHC002
Empty
D
C
ns
shHSPA8
LAMP-2A
ns
LAMP-2A
HSPA8
Total protein
CD11b^+^ cells (%)
-
+
shHSPA8
CTRL
LAMP-2A
Day4
Day2
Day4
Day2
DMSO
DMSO
ATRA
ATRA
ATRA
E
CTRL
Day4
Day2
DMSO
LAMP-2A
CTRL
NBT positive cells (%)
ns
LAMP-2A
Day4
Day2
DMSO
ATRA

### Slide 5
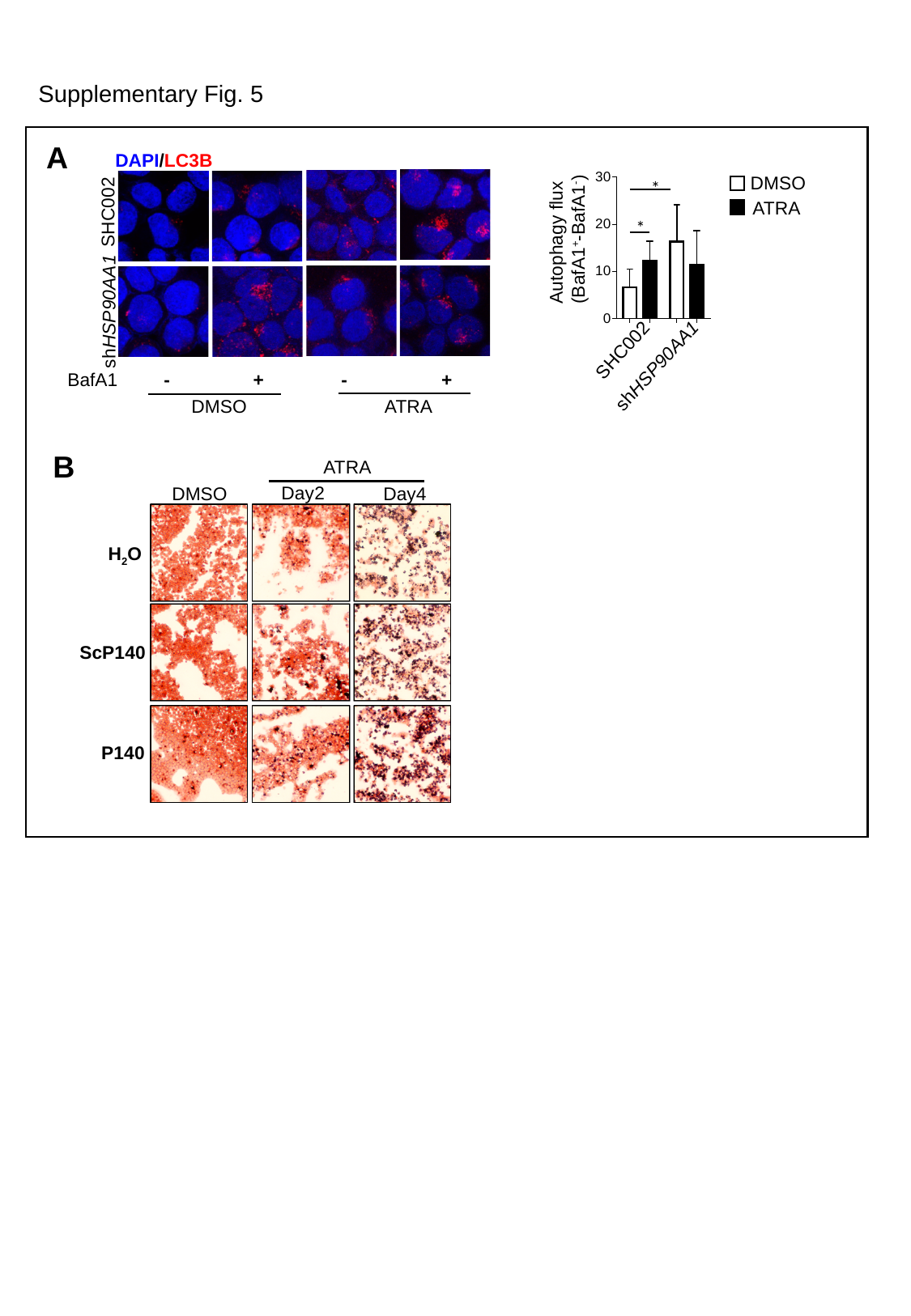

Supplementary Fig. 5
A
DAPI/LC3B
DMSO
*
ATRA
SHC002
*
Autophagy flux
(BafA1^+^-BafA1^-^)
shHSP90AA1
SHC002
shHSP90AA1
BafA1
-
+
-
+
ATRA
DMSO
B
ATRA
Day2
DMSO
Day4
H_2_O
ScP140
P140
